# Temperature, phenology, and plant defenses predict fitness near colder range limit

**DOI:** 10.1101/2023.09.11.557202

**Authors:** Daniel N. Anstett

**Affiliations:** Plant Resilience Institute, Michigan State University, East Lansing, MI 48824 USA; Department of Plant Biology, Michigan State University, East Lansing, MI 48824 USA; Department of Entomology, Michigan State University, East Lansing, MI 48824 USA; Program in Ecology, Evolution, and Behavior, Michigan State University, East Lansing, MI 48824 USA; Biodiversity Research Centre and Department of Botany, University of British Columbia, Vancouver, British Columbia V6T 1Z4, Canada

**Keywords:** Local adaptation, climate gradient, seed production, flowering time, herbivory, ellagitannins, *Oenothera bennis*

## Abstract

The space for time substitution posits that warmer locations can provide a source of genetic variation that could be adaptive for future climate change conditions. While this approximation might be useful for planning assisted gene flow, it relies on the importance of abiotic adaptations over biotic ones. Here I address this gap by assessing influence of anti-herbivore defenses, phenology, and morphology on the seed production of 146 populations of *Oenothera biennis* close to the plant’s cold range limit. Genotypes from 2.1° South of the common garden produce more seeds than most northern lineages. Adaptations across space are a suitable substitute for climate change, but there is still substantial fitness variability. These differences were best explained by bolt date, flowering time, and greater defenses against herbivores. Given the impacts of climate change, plant defenses might already be of similar adaptive importance to phenology close to northern rage limits.

## Introduction

Climate change is exposing populations to climate extremes that are increasingly further from historical averages (IPCC 2014; Pörtner *et al*. 2022). These deviations from baseline climatic conditions are overwhelming abiotic and possibly biotic adaptations (Etterson 2004; Anderson *et al*. 2012; Franks *et al*. 2014) leading to mortality (Van Mantgem & Stephenson 2007), population decline, (Willis *et al*. 2008) and range contractions (Freeman *et al*. 2018). Many populations that were locally adapted to their abiotic environment prior to the onset of climate change (Leimu & Fischer 2008; Sanford & Kelly 2011) are becoming maladapted (Aitken *et al*. 2008; Wang *et al*. 2010; Wilczek *et al*. 2014; Anderson & Wadgymar 2020; Bontrager *et al*. 2020). These populations may have to undergo range shifts to more suitable conditions to survive (Parmesan & Yohe 2003; Parmesan 2006; Poloczanska *et al*. 2013; Fadrique *et al*. 2018), or rapidly evolve to reverse demographic decline (Bell & Gonzalez 2011; Bell 2017). Such rapid evolution can occur from standing genetic variation (Barrett & Schluter 2008), yet extreme climate change may require adaptations beyond locally available genetic variation.

The space-for-time substitution is a useful approximation in which spatial biodiversity patterns across abiotic gradients are used to predict future temporal changes (Pickett 1989; Blois *et al*. 2013a). Space-for-time has been used widely to study community composition (Walker *et al*. 2010; Blois *et al*. 2013a), phenology (Buyantuyev *et al*. 2012), range-shifts (Eskildsen *et al*. 2013), and landscape-level climate maladaptation (Fitzpatrick & Keller 2015; Exposito-Alonso *et al*. 2018; Dauphin *et al*. 2021). When invoking space-for-time in the context of climate change adaptation, missing adaptive genetic variation from higher latitudes are assumed to be present in warmer populations, such that space can substitute for adaptations needed across time (Wogan & Wang 2018). Indeed, climate change experiments suggest that genetic variation from warmer localities can perform better in historically colder locations (Anderson & Wadgymar 2020; Bontrager *et al*. 2020). However, space-for-time fails to consider non-analogue future climates (Williams & Jackson 2007), unchanging landscape features such as photoperiod (Walker *et al*. 2019), and increasingly strong biotic interactions as higher latitude sites become warmer (Blois *et al*. 2013b; HilleRisLambers *et al*. 2013).

How the space-for-time substitution is influenced by differences in biotic interactions between populations remains largely untested (Bucharova 2017). Plant populations may be particularly vulnerable to increases in biotic interactions such as herbivory, which have been documented to increase at higher latitudes due to greater herbivore biomass and increased intensity of herbivore outbreaks (de Sassi & Tylianakis 2012; DeLucia *et al*. 2012; Ju *et al*. 2015; Jactel *et al*. 2019). Plants have evolved chemical and physical defenses to protect against herbivory (Ehrlich & Raven 1964; Agrawal & Weber 2015) as well as herbivore avoidance strategies such as changes in phenology (Carmona *et al*. 2011). However, biogeographic theory predicts that populations at higher latitudes will have lower plant defenses (Dobzhansky 1950; MacArthur 1972; Schemske *et al*. 2009; Baskett & Schemske 2018) (but see Moles *et al*. 2011; Anstett *et al*. 2016). This could leave higher latitude populations poorly defended and vulnerable to more mobile insect herbivores and may make plant defenses adaptive during climatic change.

Nevertheless, assuming genetic variation from lower latitudes will be effective in mediating herbivory at higher latitude sites ignores the complex and variable nature of plant-herbivore interactions. Latitudinal patterns in herbivory are notoriously variable across species (Moles *et al*. 2011; Anstett *et al*. 2016), meaning that some systems might naturally have higher herbivory in colder climates. Herbivore identity and importance can also vary latitudinally with differences in herbivory varying across herbivore guild and specialization (Hiura & Nakamura 2013; Anstett *et al*. 2014). In these conditions, plant defenses that protect against one herbivore species might be less useful on another (Krischik *et al*. 1991; Kursar *et al*. 2009; Ali & Agrawal 2012) or could even act as a feeding stimulant (Malcolm & Brower 1989; Holeski *et al*. 2013; Singh *et al*. 2021). Thus, genetic variation from donor sites could introduce plant defense traits which could be unhelpful, or even attract greater herbivore damage. Similar arguments could be made about other biotic interactions, making it vital to test the space-for-time substitution using common gardens that measure biotic interactions and traits that mediate them.

A common garden can establish what historical climate best predicts performance, and what biotic and abiotic adaptations may underly this increased performance. Multiple common garden experiments have been carried out in tree species, testing the performance of range-wide genetic variation across the range of the species including close to colder range limits (Schreiber *et al*. 2013; Aitken & Bemmels 2016; Risk *et al*. 2021). Yet less focus has been placed on testing the performance of range-wide genetic variation in forbs (but see Anderson & Wadgymar 2020) and comparing what traits provide the biotic and abiotic adaptations that cause performance differences. Given their shorter generation time, forbs might be more able to rapidly evolve and establish, yet their smaller size might make them more vulnerable to larger herbivore loads.

To address these gaps, I investigate the performance of genotypes from across the range of *Oenothera biennis* in a northern common garden, extending prior work on genetically-based clines in anti-herbivore defenses and plant phenology (Anstett *et al*. 2015). Specifically, I ask: (1) Does geography and environment of origin of populations from across the range of *O. biennis* predict seed production when grown close to the northern range limit? (2) Does increased herbivory predict decreased plant performance? (3) What phenological, morphological, and defensive traits predict performance? This approach establishes that plant defenses are as important as abiotic adaptations in predicting plant performance at higher latitudes and in explaining the traits underlying space-for-time substitution.

## Methods

### Study system

Common evening primrose, *Oenothera biennis* L., Onagraceae, is a facultatively biennial, semelparous species found along disturbed habitats such as roadsides and along bodies of water (Dietrich *et al*. 1997). This herbaceous forb is found throughout Eastern North America with limited populations in Western North America (Fig. 1), spanning a subtropical to temperate climate gradient. Due to permanent translocation heterozygosity (PTH), *O. biennis* is functionally asexual (Cleland 1972) and has near 100% selfing (Maron *et al*. 2018; Johnson and Godfrey in review). Thus populations are usually dominated by a clonal single genotype, and produce genetically identical seeds (Levin 1975). This mating system in conjunction with a semelparous life history make it possible to measure total lifetime male and female fitness in terms of the number of seeds produced by each plant.

**Figure 1.**
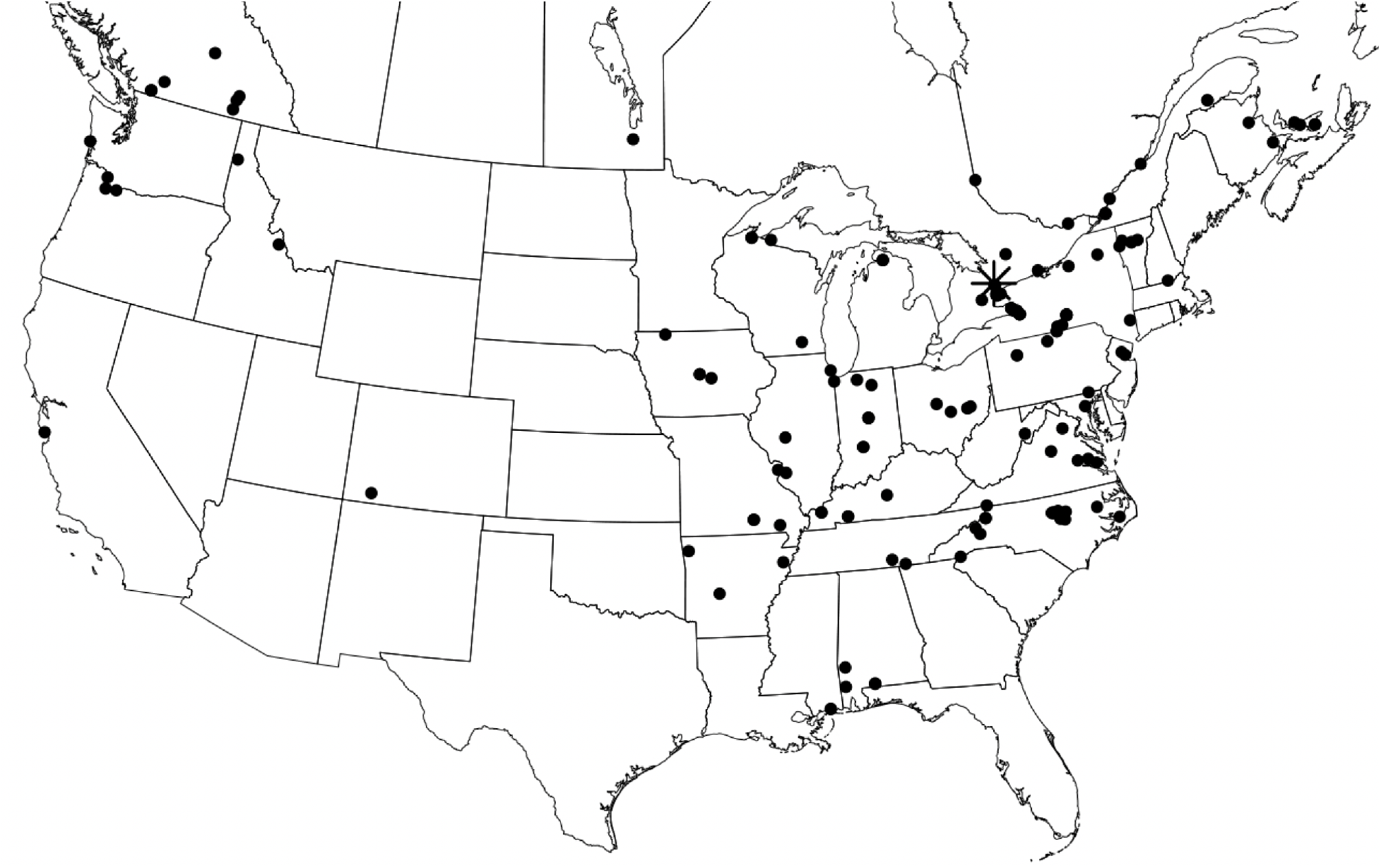
Locations of 146 sampled populations of *O. biennis* across North America. Each black dot represents one sampled population. The star represents the Koffler Scientific Reserve, the location of the common garden.

*Oenothera biennis* is an interesting case study for considering biotic adaptation during climate change. It has substantial heritable genetic variation in morphological, physiological, life history and herbivore resistance traits (Johnson *et al*. 2009) and established latitudinal gradients across many of these traits (Anstett *et al*. 2015). A diverse community of herbivores attacks *O. biennis*, including chewing herbivores, xylem feeders, and multiple specialized seed predators (Johnson & Agrawal 2005). Attacks by seed predators target the plant’s direct link to seed production and fitness and have been shown to drive rapid evolution (Agrawal *et al*. 2012). These seed predators are known to impact plants more at higher latitudes, while leaf chewing herbivory shows no association with latitude (Anstett *et al*. 2014). Phenolics and specifically the ellagitannin compound oenothein A have been associated with decreased seed predation and increased seed production (Johnson *et al*. 2009; Agrawal *et al*. 2012). Total phenolics from *O. biennis* and a mixture of oenothein A and structurally similar polymers decrease survival of both generalist and specialist insect herbivores of *O. biennis* (Anstett *et al*. 2019). Ellagitannins such as oenothein A are highly reactive in alkaline conditions (Salminen & Karonen 2011), such as those found in lepidopteran insect guts (Appel 1993), making this compound valuable to anti-herbivore defense.

### Experimental design

Seeds from 146 populations of *O. biennis,* including 132 from the main range and 14 from Western North America (Fig. 1), were sourced from established seed collections provided by Marc T.J. Johnson. These seed lines were collected one to three decades before the start of this experiment and kept in dry conditions in a -20 freezer until germination. Five replicates per population were germinated and grown for three weeks, and then planted in a common garden at the Koffler Scientific Reserve (New Market, Ontario, Canada) and grown during 2012 and 2013 as outlined in Anstett *et al*. (2015). Because of the PTH genetic system, these replicates are a single clone from each of the 146 populations. I previously characterized latitudinal gradients in physical and chemical defense traits, resistance to local herbivores and phenological differences across the range of *O. biennis*. Here I assess how geography and climate of origin of *O. biennis* impacts seed production, and what traits explain performance when measured as total lifetime fitness. The timing of the common garden captures the early effects of climate change in North America with mean annual temperature being 1.6 °C warmer at the common garden across the 2012 and 2013 period when compared to the historical baseline temperatures under which *O. biennis* evolved.

### Climate & geography

Historical climate data from 1960 to 1990 was downloaded from Climate NA for each population (Wang *et al*. 2016), which is representative of pre-climate change conditions. I selected seven temperature and four precipitation associated variables that could impact *O. biennis* fitness. Most variables were highly correlated (r > 0.7; Table S1), hence only mean annual temperature (MAT), mean summer precipitation (MSP), cumulative moisture deficit (CMD), and relative humidity (RH) were included in downstream calculations. Monthly climate data was downloaded from Climate NA for 2012 and 2013 for the Koffler Science Reserve (44.026479, -79.544686). MAT, CMD, and RH was calculated as a combined metric for July 2012 to September 2013, by first averaging July, August and September values across both years and then calculating a 12 month mean. This calculation reflects the entire length of the common garden experiment. MSP for the common garden was only included for 2013 since supplemental watering was provided in 2012 after seedling transplant. Climatic distance to the common garden was calculated for each climatic variable by subtracting the common garden value from the 30-year mean.

### Trait measurement

I measured 23 traits including phenological (flowering date, bolt date, growth rate); morphological (leaf toughness, trichome number, specific leaf area (SLA); % water content), secondary chemistry across leaf, flower and fruit tissues (total phenolics, oxidative capacity; concentration of ellagitannin compounds oenothein A and oenothein B); and herbivore resistance traits (generalist leaf herbivory, number of *Philaenus spumarius* L. Hemiptera: Aphrophoridae, fruits damaged by specialist *Schinia florida* G. Lepidoptera: Noctuidae, fruits damaged by specialist *Mompha brevivitella* C. Lepidoptera: Momphidae). The detailed methodology for the estimation of 22 of these traits is found in Anstett *et al*. (2015). Additionally, I monitored bolt date throughout the season, defined as the first day the stem length was greater than 5 cm. The total number of fruits and the length and width of 5 fruits produced by each plant was recorded. Seed number was estimated by the following formula: seeds number = 2.33*Average_Fruit_Length * Average_Fruit_Width - 102.47 * Damaged_Fruits. This calculation has been established to accurately estimate fruit number in prior work conducted on *O. biennis* (Agrawal *et al*. 2012). The damaging effects of specialist seed predators were also taken into account in the final estimation of seed production by subtracting the total number of *S. florida* and 0.18\**M. brevivitella* damaged fruits. The latter represents average known amount of seed loss per pod due to *M. brevivitella* damage (M.T.J. Johnson personal communication).

### Statistical analyses

All analyses were performed using R (R Development Core Team 2020). Means of each population were used for all traits ensuring a comparable design across all traits, since dried, ground plant issue was averaged across genotypes prior to chemical analysis. I predicted seed number using geographic, climatic, herbivory and plant traits. Seed number had inflated zeros and was overdispersed (P <0.001) using *dispertiontest* in the *AER* package in R (Kleiber *et al*. 2020). Hence, all regressions predicting seed number were performed using a negative binomial generalized linear model in the *MASS* package in R (Ripley *et al*. 2013). For each model, I tested first order patterns, and quadratic patterns (if this pattern appeared possible in the raw data) (Table S2). Quadratic predictors were assessed separately from first order predictors using the I(Trait^2) function in R. Regression models were reduced by using stepwise AIC, selecting the model with the lowest AIC score. Significance of this model was assessed using a type 2 anova using the *Anova* command in the CAR package in R (Fox *et al*. 2012). Results were visualized using visreg in the *visreg* package in R (Breheny *et al*. 2020), with scale= “response”, which applies an inverse log-link function scaling the plots in seed number units.

Seed number was modeled as a function of latitude and longitude, distance to common garden, climate present at each population, and climate distance to the common garden (Table S2). Then to determine what types of traits underly the geographical and climatic patterns, resistance to herbivores, phenology, morphology, and plant secondary chemistry were each used to predict seed number. I used multiple regression within each class of traits whenever possible (see Table S2 for every tested model). Leaf toughness was highly correlated with other morphological variables (r>0.7) and was not included in the morphology multiple regression. For resistance to herbivores, I ran separate regressions for seed predators. For secondary compounds, I ran regressions separately for each tissue. Total phenolics was assessed separately from individual compounds. Oxidative capacity was highly correlated with total phenolics (r>0.7); oenothein B was highly correlated with oenothein A. Hence, I selected only total phenolics and oenothein A based on prior knowledge of what traits more strongly predict resistance to herbivores (Agrawal *et al*. 2012; Anstett *et al*. 2015).

To understand the major axes of variation across these predictive variables, I also carried out principal component analysis using *prcomp()* in R and associated the most important axes with seed production. Biplots were visualized using *autoplot()* in the ggfortify package in R (Tang *et al*. 2016). To further establish which variables are most important in predicting *O. biennis* seed number, I compared the relative importance of all variables that had evidence of predictive seed number (flowering date, bolt date, water content, growth rate, and number of trichomes as well as leaf, flower and fruit total phenolics and oenothein A). I carried out this comparison with the machine learning algorithm, *randomFores*t in R (Liaw & Wiener 2002) using 1000 trees and plotted relative variable importance. I further visualized the relationship between latitude, seed number and six of the most important traits by plotting trait change over latitude, while color coding points for seed number using ggplot2 in R (Wickham 2011).

All analyses were repeated using undamaged fruits as a response instead of the estimated seed number. No major differences were detected in trends and only minor difference in significance (data not shown); hence I present only the seed number results.

## Results

### Geography & climate

Seed production increased with latitude, peaking at 41.9° N, 2.1° south of the common garden before declining (Fig. 2A, Table S3; linear P = 0.001, quadratic P = 0.001). Indeed, five of the top six genotypes with the most fitness were from 41.9°N or further south. Models with longitude had higher AIC and were not selected. Seed production decreased at distances further from the common garden (P = 0.03; Fig. 2B), although the highest performing genotypes were from 100’s or 1000’s of km from the common garden.

**Figure 2.**
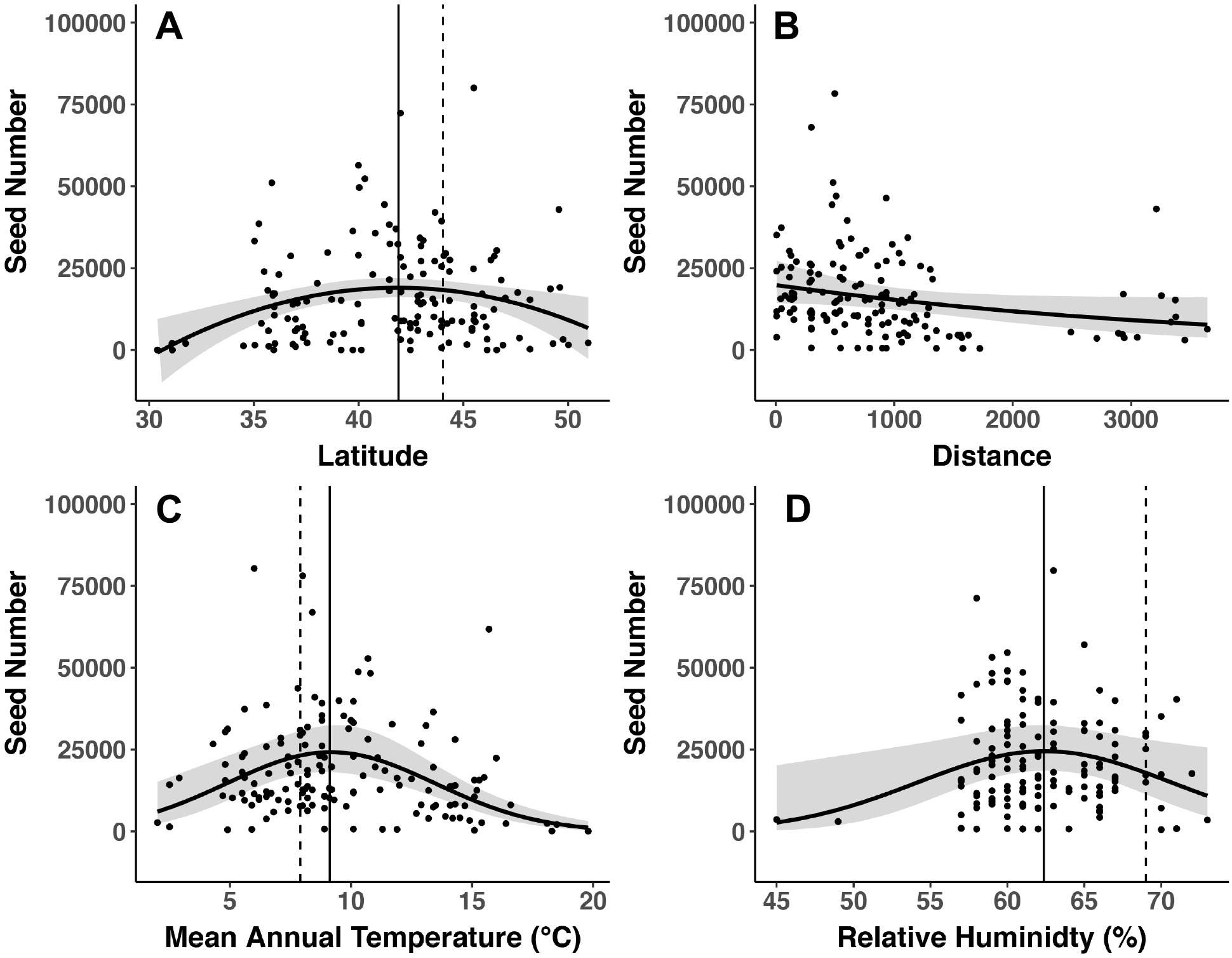
Seed production of *O. biennis* in a Northern common garden as predicted by (A) latitude of origin, (B) Euclidian distance to the common garden in km, (C) mean annual temperature at collected location, and (D) relative humidity at collected location. The solid vertical line gives the latitude or environmental condition at origin of population with the highest predicted seed production. The dashed vertical line gives the latitude or environmental condition at the common garden location. Each point represents one population.

Temperature and relative humidity explained plant fitness. Seed production increased towards genotypes from warmer temperatures, peaking at 9.1°C, 1.2°C warmer than the common garden before declining at higher temperatures of origin (Fig. 2C; linear P = 0.004, quadratic P = 0.001). Genotypes from locations with greater relative humidity had higher seed production up to 62.4% relative humidity, 6.6% higher than the common garden, before declining at higher relative humidity (Fig. 2D). However, this effect was only weakly supported (linear P = 0.057, quadratic P = 0.055). Models with May to September precipitation and Hargreaves climatic moisture deficit had higher AIC and were not selected. Temperature distance from common garden has a similar effect with genotypes from locations with greater temperature distance peaking 1.4°C warmer from the common garden before declining (Fig. S1, Table S3, P < 0.002). There was also weak support for mean summer precipitation distance predicting seed number (P = 0.083; Table S3). Models with relative humidity distance and Hargreaves climatic moisture deficit distance were not selected.

### Resistance to herbivory

Herbivory had variable relationships with seed production (Table S3). Seed production increased with *Philaenus spumarius* number of to 14 individuals and then declined (Fig. 3A; linear P <0.001, quadratic P = 0.01). The model that included leaf herbivory had higher AIC and was not selected. Seed predators had contrasting relationships with seed production. Seed number increased with higher *S. florida* damage or *S. florida* simply attacked more fecund plants (Fig. 3B; P = 0.003). Seed number decreased with greater *M. brevivitella* damage although there was considerable variability (P = 0.058; Fig. 3C).

**Figure 3.**
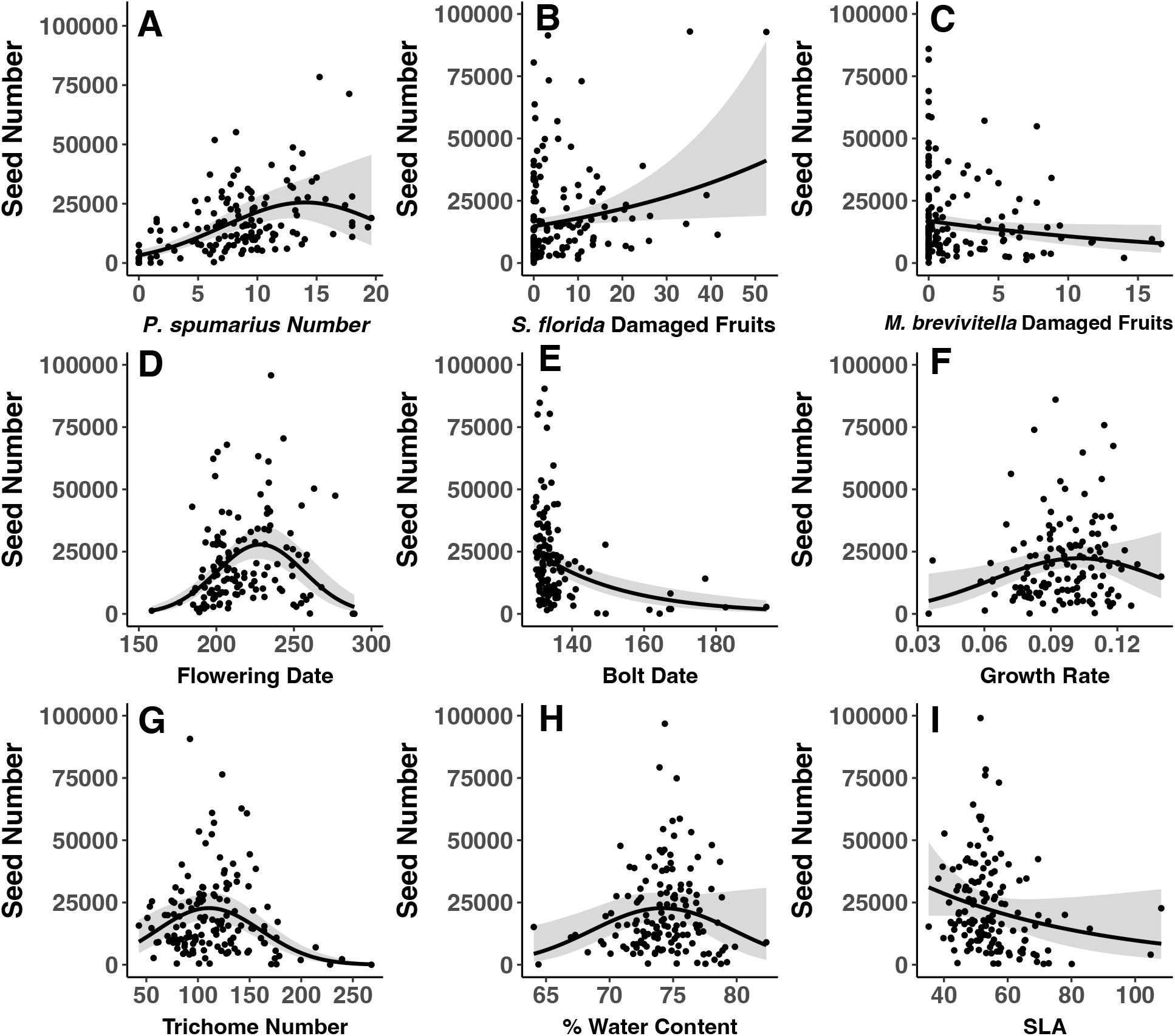
Herbivory, phenology and plant morphology predicts seed production in *O. biennis* in a northern common garden. Plots are given for (A) number of *Philaenus spumarius*, (B) number of *Schinia florida* damaged fruits, (C) number of *Mompha brevivitella* damaged fruits, (D) flowering date (in Julian days), (E) bolt date (in Julian days), (F) growth rate (cm/d), (G) trichome number in a 5 mm diameter disk, (H) % water content, and (I) specific leaf area (SLA). Each point represents one population.

### Plant phenotype

All phenology variables predicted *O. biennis* seed number in some way (Table 3). Plants with early or late flowering date performed poorly, with peak flowering date occurring at an intermediate flowering date (Fig. 3D; P < 0.001). Genotypes with later bolt date had low seed production, while plants with earlier bolt date showed higher seed production (on average) but with higher variability (Fig. 3E; P < 0.001). Plants with intermediate growth rate had higher seed production although this relationship was weak (linear P = 0.049, quadratic P = 0.071; Fig. 4F). Most plant morphology variables also predicted seed number (Table S3). Seed production increased with trichome number up to 110 per 5 mm^2^, before decreasing at higher trichome numbers (Fig. 3G; P < 0.001). Fitness also peaked at intermediate water content (P = 0.03), with seeds increasing with water content up to 74.3% before decreasing (Fig. 3H). Finally, seed number was not associated with SLA (Fig. 3I; P = 0.11).

**Figure 4.**
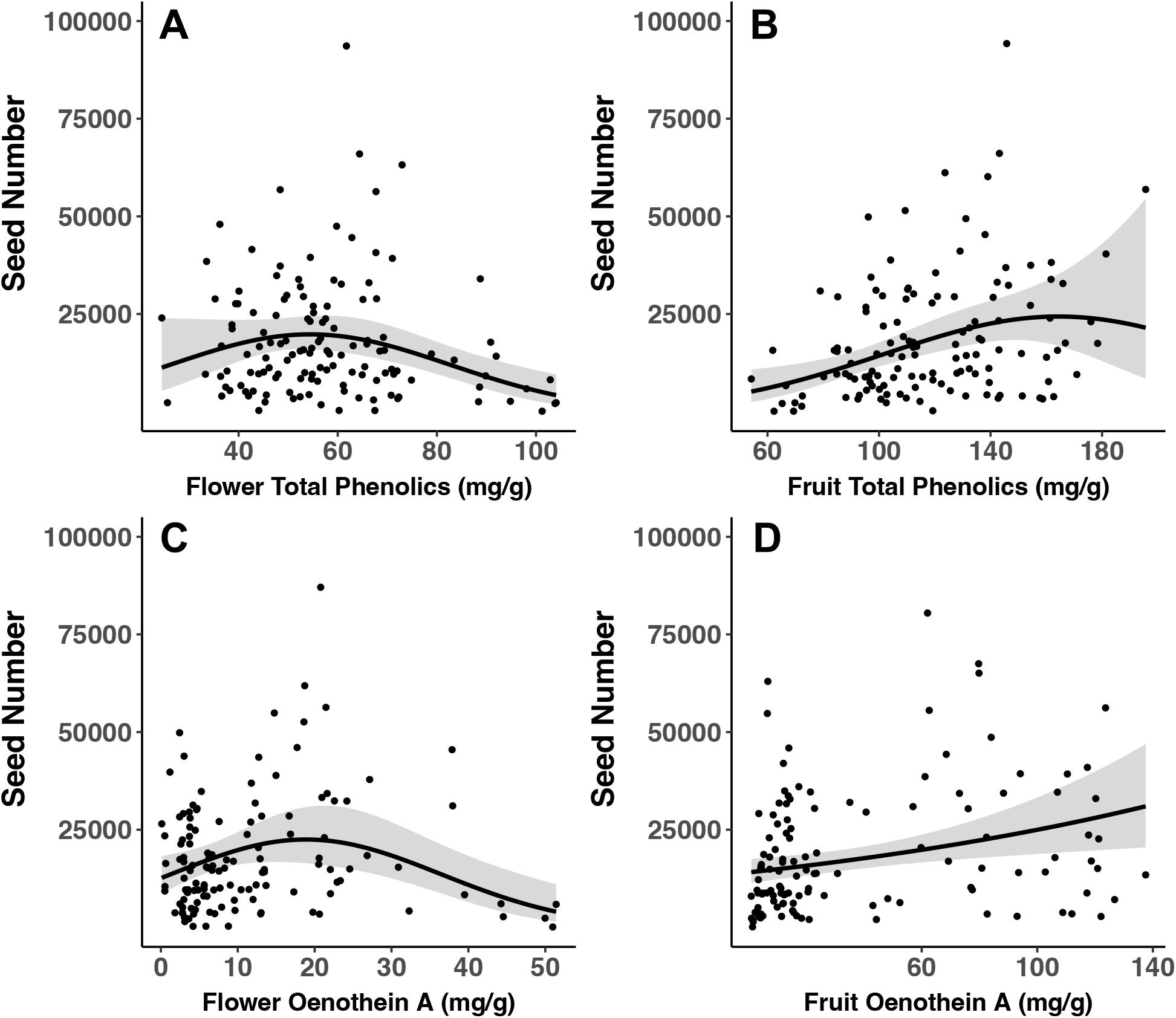
Fruit chemical defenses predict seed production of *O. biennis* in a common garden. Plots are given for (A) flower total phenolics, (B) fruit total phenolics, (C) flower oenothein A, and (D) fruit oenothein A. Units are in mg of given secondary metabolite(s) per g of dry mass. Each point represents one population.

Plant defense chemistry metrics also predicted seed production but only for some tissues (Table S4). Higher leaf total phenolics weakly predicted greater seed production (Fig. S2A; P = 0.069), while leaf total oenothein A does not predict seed number (linear P = 0.44; quadratic P = 0.76; Fig. S2B). Higher flower total phenolics predicted greater seed number up to 54.7 mg/g and then declined (Fig. 4A; linear P = 0.045, quadratic P =0.02). Seed number increased with flower oenothein A (Fig. 4C; linear P = 0.01) and declined slightly at higher values (quadratic P = 0.01). Greater fruit total phenolics predicted greater seed production (Fig. 4B; P = 0.03) with poor evidence of decline at high fruit total phenolics (P = 0.12). Seed number also increases with greater fruit oenothein A (P = 0.006; Fig. 4D).

Principle component analyses revealed that most of *O. biennis* trait variation was predicted by the first two principal components (Fig. S3; Table S5), with PC1 explaining 33.65% of the variation and PC2 explaining 13.8%. PC1 is primarily associated with flowering date, fruit oenothein A, fruit phenolics, flower oenothein A, leaf oenothein A, leaf phenolics, and bolt date, while PC2 is primarily associated with growth rate, SLA, and water content (Fig. 5A, Table S6). Smaller values of PCA 1, associated with later phenology and higher plant defenses predicted greater seed number (P=0.02; Table S4). Smaller values of PC2, associated with decreased SLA, growth, rate, and water content predicted greater seed number (P=0.004), but with much variation and with the highest seed producers having intermediate seed numbers.

**Figure 5.**
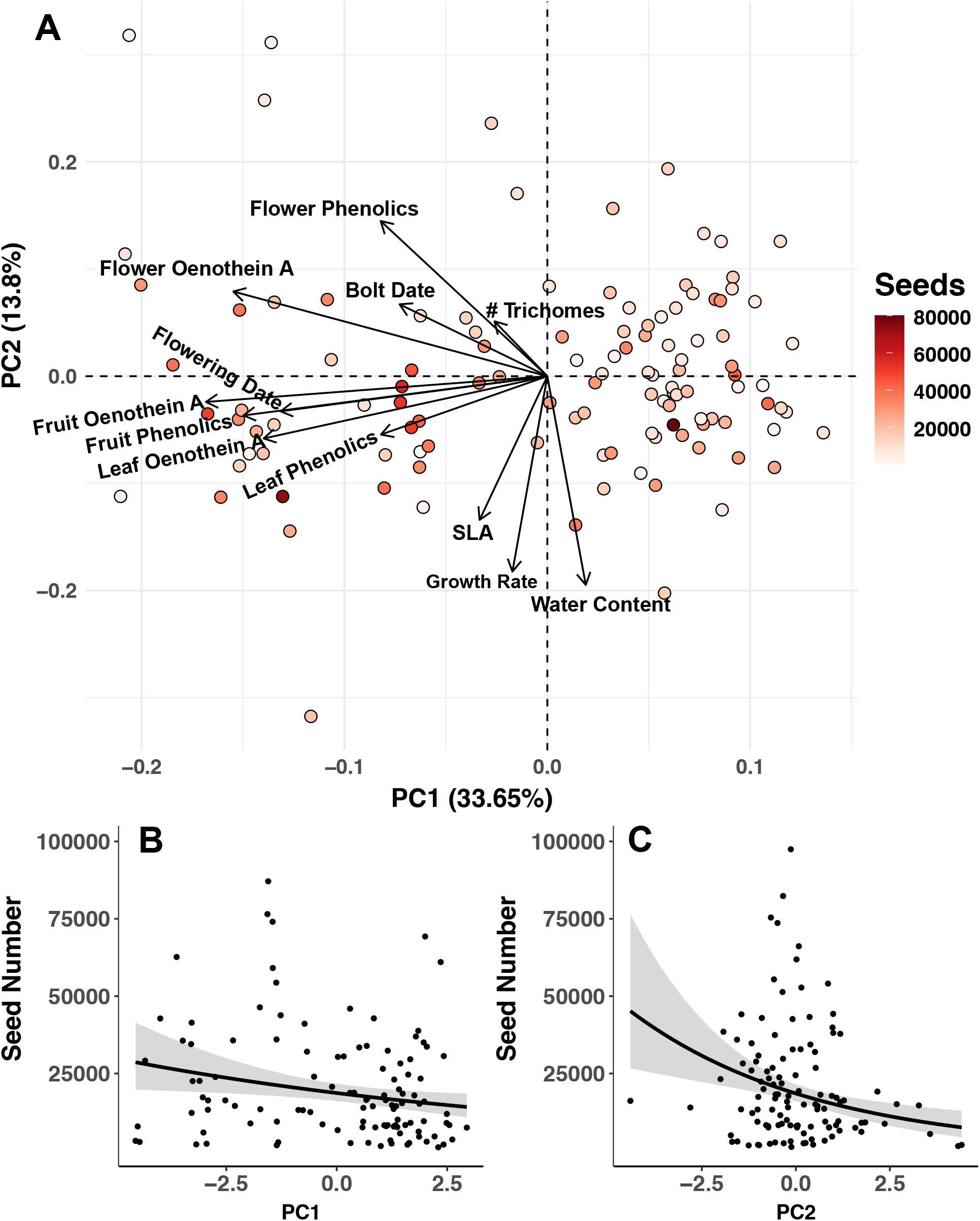
Principal component analyses of chemical, phenological and morphological traits of *O. biennis*. (A) PCA bioplot showing first and second principal component. PC1 describes mostly chemical and phenological traits, while PC2 describes primarily morphological traits and growth rate. Each dot represents one population. Number of seeds per plant is color coded for each point. Seeds are predicted by (A) PC1 and (B) PC2. Each point represents one population.

Gradient forest analysis showed that, out of the variables predictive of seed number, bolt date, flower oenothein A, and fruit oenothein A were the most important (Fig. 6). Fruit total phenolics, and flowering date were next in variable importance. Then growth rate, flower total phenolics, water content, and trichome number were the least important. Latitude-trait plots with color-coded seed production emphasized flowering time just south of the common garden leads to increased seed production (Fig. S4A), while later bolt date from further south leads to massively decreased seed production (Fig. S4B). Populations from directly south of the common garden that have greater fruit defenses perform considerably better (Fig. S5). Flower defenses south of the common garden are also somewhat predictive of greater seed production (Fig. S6).

**Figure 6.**
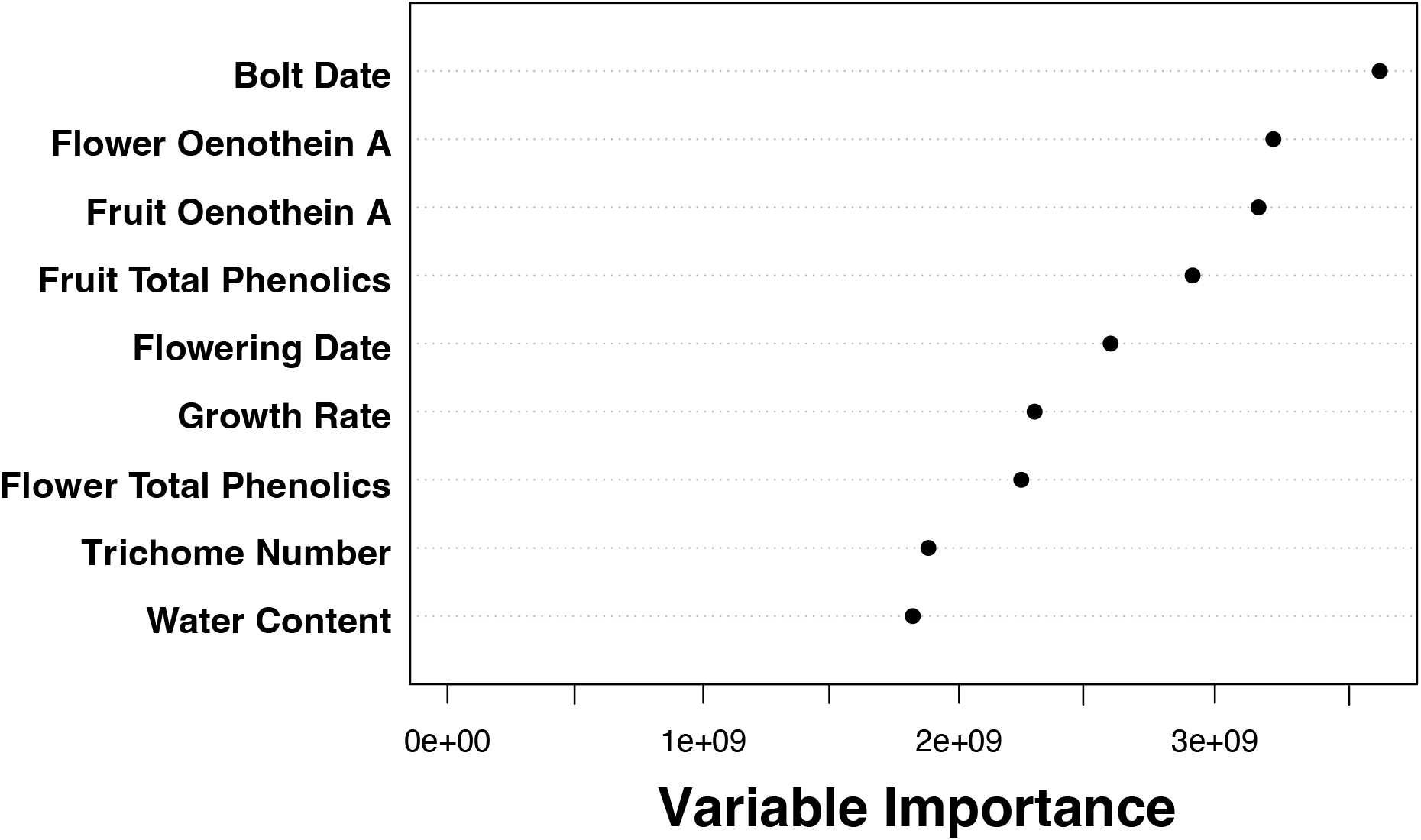
Variable importance for *O. biennis* traits predicting seed production in a Northern common garden. Values were generated using gradient forest.

## Discussion

This study shows that anti-herbivore defenses can at least partially explain the space-for-time substitution that genotypes from warmer latitudes are more adapted to climate change conditions. A 1.6 degree increase from historical baseline temperatures has made *O. biennis* genotypes from 2.1° South of the common garden perform better than local more northern genotypes. This performance difference is best explained by both phenology and plant phenolics, suggesting both biotic and abiotic factors impact performance during the early effects of climate change. The importance of phenology as a climate change adaptation is already well established (Hamann *et al*. 2018; Wadgymar *et al*. 2018). Here I extend our understanding of climate adaptation by implicating the benefit of increased plant chemical traits typically associated with plant defense as an important part of climate change adaptation.

### Climate change adaptation close to cooler range limit

Plant defenses can be an important adaptation at higher latitude sites and could even be more important than abiotic adaptations. Here I show that genotypes from across the range of *O. biennis* with greater fruit total phenolics, greater fruit oenothein A and intermediate values of flower oenothein A (mostly higher than the local population) had greater seed production when grown close to the cold range limit of *O. biennis* (Fig. 4-5, Fig. S5-6). Additionally, *O. biennis* chemical traits ranked higher in variable importance than flowering time (Fig. 6), which is usually more commonly associated with the impacts of climate change. This importance of plant defenses at higher latitudes in *O. biennis* is supported by prior work showing that greater oenothein A explains decreased seed predator damage in this common garden (Fig. S4A; Anstett *et al*. 2015) and during experimental evolution (Agrawal *et al*. 2012). Seed predator damage, especially by *M. brevivitella*, is also known to lead to decreased seed production (Fig. 3C; Agrawal *et al*. 2012), pointing to the importance of greater plant defenses at higher latitudes in this system.

Flowering time and bolt date were also both predictive of greater seed production, with earlier bolt date and an intermediate date of first flower being optimal. Some plants failed to bolt until much later in the season, thus producing many fewer seeds (Fig. S3B) and making bolt date the most important predictor of success for this small subset of populations (Fig. 6).

*Oenothera biennis* populations from the southern part of the range of the plant showed a suboptimal, much later flowering time when grown in the northern common garden (Anstett *et al*. 2015) likely due to photoperiod and/or temperature cue issues, which are often raised when considering phenotypic mismatch during latitudinal translocations (Wadgymar & Weis 2017).

Indeed, these plants would have less time to acquire resources and produce seeds before the end of the growing season (Inouye 2008; Franks 2015). Thus, plants from higher latitudes that are just South of the common garden were better able to produce more seeds because they likely had a photoperiod closer to the garden.

This begs the question as to why an intermediate flowering date leads to greater seed production in *O. biennis* rather than the earliest possible flowering date. One possibility is that these early flowering plants developed too early and cooler weather damaged the plants. Another explanation is that plants that flower earlier are available for longer in the season and are thus more susceptible to herbivory. Indeed, prior work in this system suggests that genotypes with earlier flowering have greater damage by generalist and specialist herbivores (Agrawal *et al*. 2013; Anstett *et al*. 2015). This pattern is not surprising given that life history traits have been found to be consistent predictors of resistance to herbivores (Pilson 2000; Kawagoe & Kudoh 2010), and at times more reliable than traditional chemical or physical defenses (Carmona *et al*. 2011). Overall, this suggests that even phenological adaptations at northern range limits may be adaptations to biotic interactions.

### Implications for space-time substitution

Using space as a proxy for time during translocations takes into account the effects of temperature adaptation on average, but ignores how genetics within a region may impact the realized fitness of a randomly selected individual. On average, genotypes from south of the common garden with slightly higher mean annual temperature perform better (Fig. 2A,C), while other climate traits were less important. This matches previous findings pointing to temperature as a key factor in field transplant experiments (Bontrager *et al*. 2020). However, average temperature or latitude may hide considerable genetic variability within a particular temperature band. Multiple genotypes from latitudes at or slightly above the common garden still had above average seed number (Fig. 2A), while some genotypes from across the entire range had low fitness. This variability points to the importance of testing on-the-ground fitness at future introduction sites or at least considering the variety of abiotic and biotic adaptations that might be needed and where this genetic variation might be sourced.

This system suggests how possible trade-offs between biotic and abiotic adaptation may be an important consideration for assisted gene flow. The benefits of assisted gene flow as a way to rescue populations from climate change has frequently been weighed against risks of introducing maladapted genetic variation (Vitt *et al*. 2010; Aitken & Whitlock 2013). Space-for-time substitutions might serve as a way to source genetic material for assisted gene flow while mitigating the risks of introducing further maladaptation in an already vulnerable species. This study suggests there are risks of introducing genotypes that may be well suited for the new abiotic environment but not the biotic one (or vice versa). These trade-offs are possible in *O. biennis* especially given that multiple traits explain fitness and that these types of traits are known to covary (Johnson *et al*. 2009). Given these findings, studies focusing on species of immediate conservation concern should investigate the possible greater importance of biotic adaptations close to northern range limits and how these may interact with abiotic adaptation (Blois *et al*. 2013b; HilleRisLambers *et al*. 2013). This may be particularly true for translocations across latitudinal gradients (Fielding *et al*. 1999), where adaptation to photoperiod may lead to poor match with phenology (Walker *et al*. 2019).

### Implications for biogeographic hypotheses

Greater plant defenses at higher latitudes may provide needed adaptations to increased herbivory. Herbivore outbreaks at higher latitude sites are being attributed directly or indirectly to climate change (de Sassi & Tylianakis 2012; DeLucia *et al*. 2012; Ju *et al*. 2015; Jactel *et al*. 2019), raising the possibility that selection for greater defenses at higher latitudes might already be occurring. Indeed, I previously documented that damage by seed predators is greater at higher latitudes in *O. biennis* (Anstett *et al*. 2014), possibly due to early effects of climate change. This idea falls against the established pre-climate change paradigm that greater biotic interactions at warmer, lower latitudes leads to selection for greater traits that mediate these interactions (Dobzhansky 1950; MacArthur 1972; Coley & Barone 1996; Schemske *et al*. 2009). However, selection for greater plant defenses at higher latitudes due to climate change might explain why more studies are finding greater plant defenses at higher latitudes (Moles *et al*. 2011; Anstett *et al*. 2016).

As the impacts of the Anthropocene become more acute, long-held biogeographic hypotheses may require a more complex and updated view. The causes and drivers of range limits has been a key question throughout the history of evolutionary biology as well as a driver of biogeographic patterns (Darwin 1859; MacArthur 1972; Sexton *et al*. 2009). Biotic interactions are viewed as being less important for determining colder range limits (Louthan *et al*. 2015; Paquette & Hargreaves 2021). Biotic interactions and the traits that mediate them are also expected to be less intense and important at higher latitude sites (MacArthur 1972; Coley & Aide 1991; Coley & Barone 1996; Schemske *et al*. 2009; Johnson & Rasmann 2011; Anstett *et al*. 2016). Both of these ideas share the common viewpoint of diminishing the importance of biotic interactions in colder climates. Climate change is changing this reality and is likely going to pose new biotic challenges on historically cooler sites that may lack adaptations for greater mean interaction intensity (de Sassi & Tylianakis 2012; Ju *et al*. 2015) and greater variability in that intensity (Wetzel *et al*. 2023). Greater biotic adaptations such as plant defenses against herbivores may become an increasingly important part of plant adaptation at higher latitudes leading to the weakening of latitudinal gradients in biotic interactions and the traits that mediate them. Lack of these adaptations at northern range limits could even slow plant range expansions that are so often evoked when studying climate change.

### Data Availability

All code and data for this paper is available on GitHub https://github.com/anstettd/KSR_North.

### Conflict of Interest

The author has no conflicts of interest.

## Acknowledgements

I thank C. Fitzpatrick, M. Turcotte, N. Turley, J. Hollister, C. Thomsen, R. Godfrey, A. Lee, P. Maharaj, L. Sequeira, T. Didiano, D. Filice, D. Vigmond, H. Sekhon, A. Longley, J. Anstett A. Koivuniemi, and J. Kim for field and lab support. J.-P. Salminen and J. Ahern aided in analytical chemistry. M. Johnson and S. Greiner provided seeds. M. Johnson, W. Wetzel, and A. VanWallendael also provided advice and read over a draft of the paper. J. Anstett aided in coding. Special thanks to A. Agrawal for advice on seed estimation in *O. biennis*. Funding was provided by Sigma Xi Grant-in-Aid of Research Grant, NSERC CGS-D Vanier, NSERC PDF, NSERC Banting and a MSU Plant Resilience Institute Postdoc Fellowship to D. Anstett.

## Notes

https://github.com/anstettd/KSR_North

### Competing Interest Statement

The authors have declared no competing interest.

### Summary of Updates

Minor corrections and updates to key figures and wording in the results section. All tables were also moved to supplement.

