## Supplemental Material for "Temperature, phenology, and plant defenses predict fitness near colder range limit"

**Supplementary material for: Temperature, phenology, and plant defenses predict fitness near colder range limit**

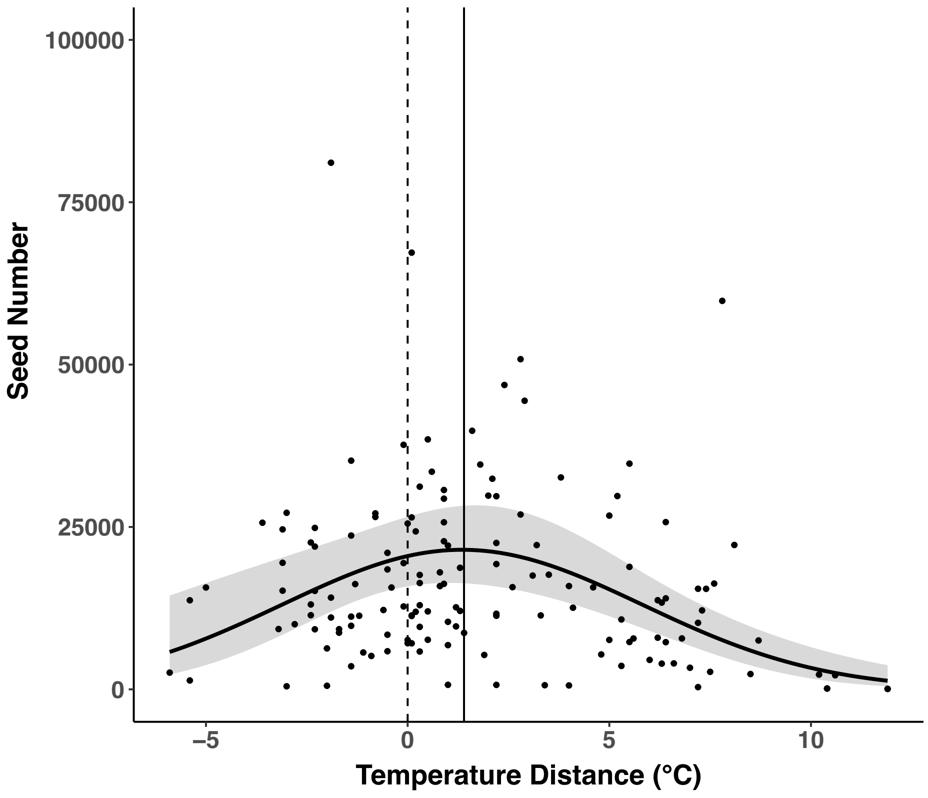

Figure S1. Seed production of *O. biennis* in a Northern common garden as predicted by temperature distance, the difference between mean annual temperature at climate of origin versus the common garden. The solid vertical line gives the environmental condition at origin of population with the highest predicted seed production. The dashed vertical line gives the environmental condition at the common garden location. Each point represents one population.

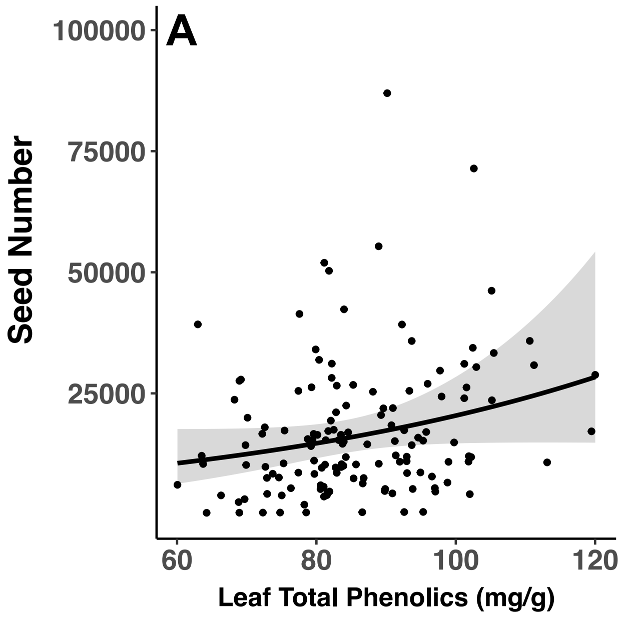

Figure S2. Seed production of *O. biennis* in a Northern common garden as predicted by (A) leaf total phenolics, and (B) leaf oenothein A. Units are in mg of given secondary metabolite(s) per g of dry mass. Each point represents one population.

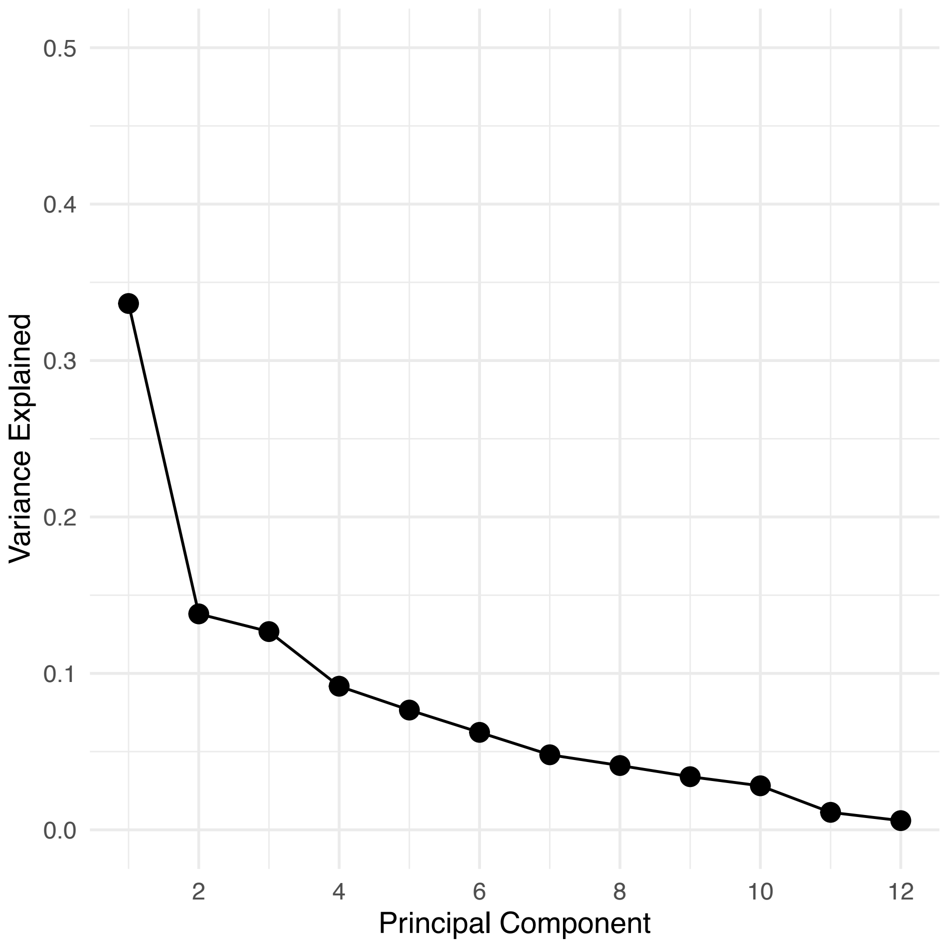

Figure S3 Scree plot showing the proportion of variance explained by 12 principal components.

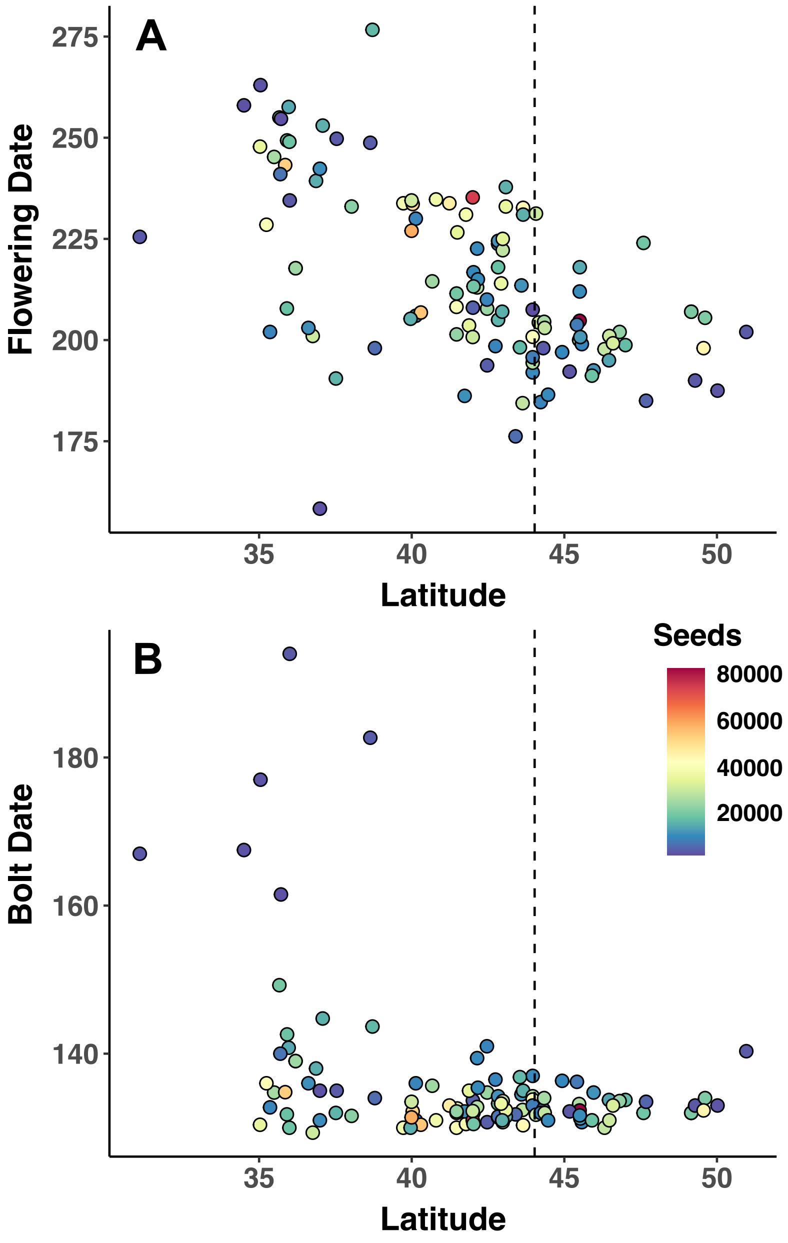

Figure S4. Genetically-based latitudinal pattern in phenology for *O. biennis*. Plots are given for (A) flowering date (in Julian days), and (B) bolt date (in Julian days). Average number of seeds produced per population is shown as a color gradient on the points with warmer colors representing increased seed production.

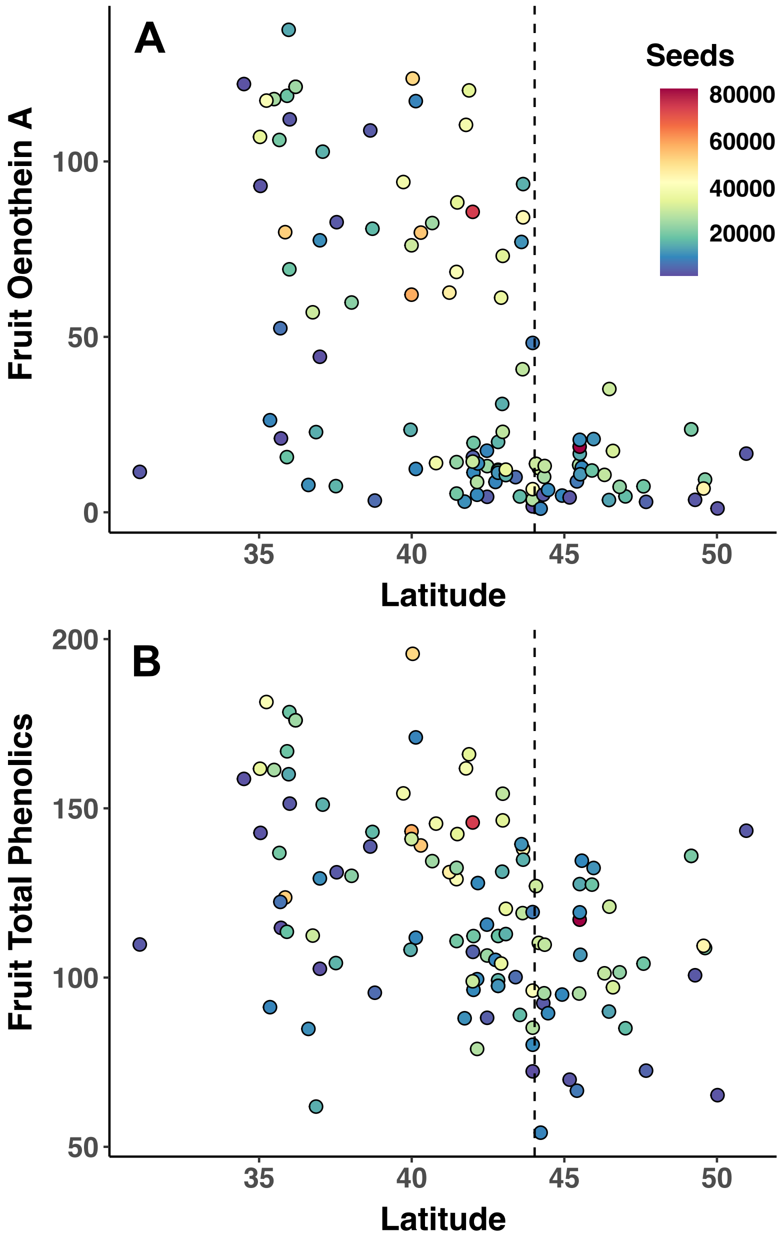

Figure S5. Genetically-based latitudinal pattern in phenology for *O. biennis*. Plots are given for (A) fruit oenothein A (mg/g), and (B) fruit total phenolics (mg/g). Average number of seeds produced per population is shown as a color gradient on the points with warmer colors representing increased seed production.

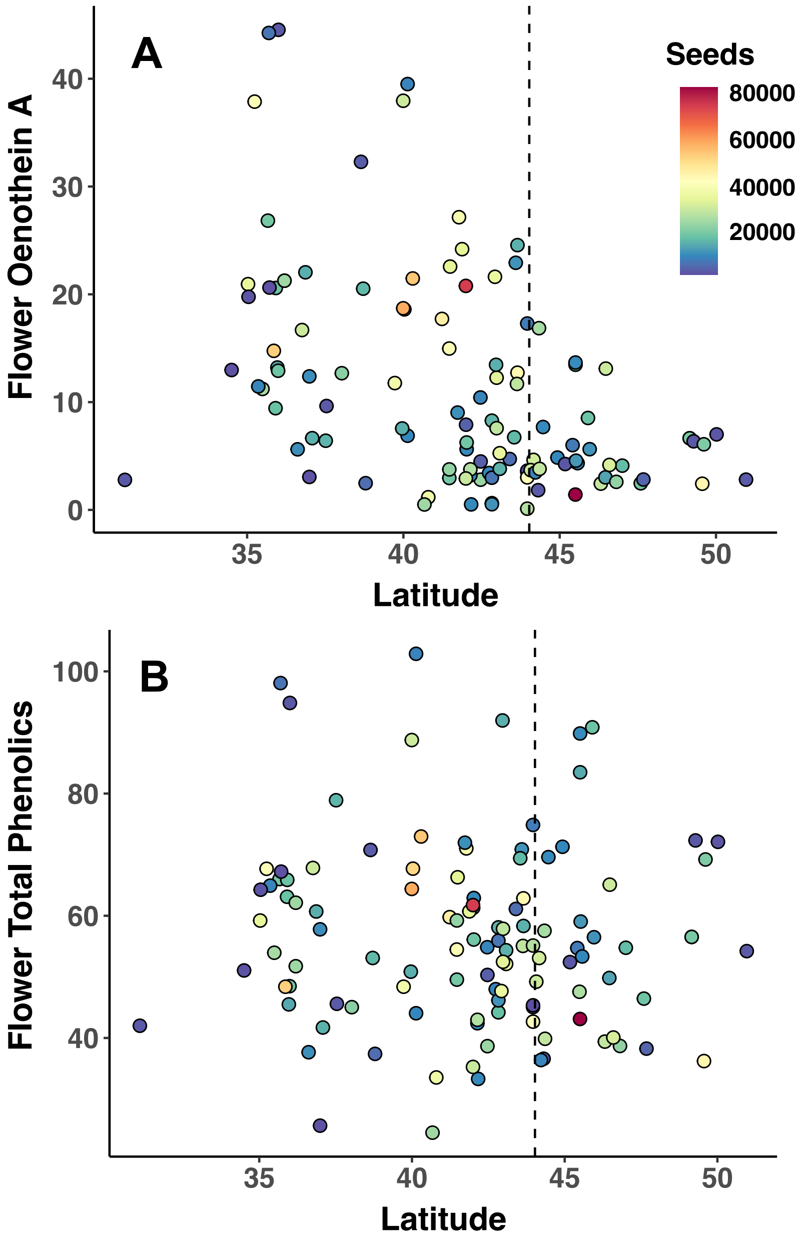

Figure S6. Genetically-based latitudinal pattern in phenology for *O. biennis*. Plots are given for (A) flower oenothein A (mg/g), and (B) flower total phenolics (mg/g). Average number of seeds produced per population is show as a color gradient on the points with warmer colors representing increased seed production.

Table S1. Pearson correlation coefficients for climate variables across 146 populations of *Oenothera biennis*.

MAT=mean annual temperature (°C), MWMT= mean warmest month temperature (°C), MCMT= mean coldest month temperature (°C), TD=temperature difference between MWMT and MCMT (°C), NFFD=number of frost-free days, DD18=degree-days above 18°C, MAR= mean annual solar radiation (MJ $m^{-2}d^{-1}$), MAP=mean annual precipitation (mm), MSP=mean summer precipitation (mm), CMD=Hargreaves climatic moisture deficit (mm), RH=mean annual relative humidity (%).

| | MAT | MWMT | MCMT | TD | NFFD | DD18 | MAR | MAP | MSP | CMD | RH |
|---|---|---|---|---|---|---|---|---|---|---|---|
| **MAT** | 1 | 0.85 | 0.94 | -0.66 | 0.86 | 0.92 | 0.75 | 0.6 | 0.5 | 0.25 | -0.04 |
| **MWMT** | 0.85 | 1 | 0.64 | -0.18 | 0.54 | 0.95 | 0.68 | 0.35 | 0.6 | 0.12 | -0.17 |
| **MCMT** | 0.94 | 0.64 | 1 | -0.87 | 0.92 | 0.77 | 0.66 | 0.65 | 0.34 | 0.31 | 0.07 |
| **TD** | -0.66 | -0.18 | -0.87 | 1 | -0.83 | -0.38 | -0.41 | -0.61 | -0.05 | -0.31 | -0.2 |
| **NFFD** | 0.86 | 0.54 | 0.92 | -0.83 | 1 | 0.71 | 0.54 | 0.61 | 0.2 | 0.32 | 0.23 |
| **DD18** | 0.92 | 0.95 | 0.77 | -0.38 | 0.71 | 1 | 0.74 | 0.51 | 0.63 | 0.12 | -0.1 |
| **MAR** | 0.75 | 0.68 | 0.66 | -0.41 | 0.54 | 0.74 | 1 | 0.46 | 0.55 | 0.14 | -0.27 |
| **MAP** | 0.6 | 0.35 | 0.65 | -0.61 | 0.61 | 0.51 | 0.46 | 1 | 0.64 | -0.36 | 0.18 |
| **MSP** | 0.5 | 0.6 | 0.34 | -0.05 | 0.2 | 0.63 | 0.55 | 0.64 | 1 | -0.59 | -0.16 |
| **CMD** | 0.25 | 0.12 | 0.31 | -0.31 | 0.32 | 0.12 | 0.14 | -0.36 | -0.59 | 1 | -0.32 |
| **RH** | -0.04 | -0.17 | 0.07 | -0.2 | 0.23 | -0.1 | -0.27 | 0.18 | -0.16 | -0.32 | 1 |

Table S2. Equations predicting seed number across 146 populations of *O. biennis*.

MAT=mean annual temperature (°C), MSP=mean summer precipitation (mm), CMD=Hargreaves climatic moisture deficit (mm), RH=mean annual relative humidity (%). SLA=specific leaf area.

| Category | Equation | Equation Selected by Stepwise AIC |
|---|---|---|
| **Geography** | Seeds = Latitude + Longitude + $Latitude^2$ + $Longitude^2$ | Seeds = Latitude + Longitude |
| **Distance to Common Garden** | Seeds = Distance | n/a |
| **Climate** | Seeds = MAT + MSP + CMD + RH + $MAT^2$ + $MSP^2$ + $CMD^2$ + $RH^2$ | Seeds = MAT + RH + $MAT^2$ + $RH^2$ |
| **Climate Distance** | Seeds = MAT Distance + MSP Distance + CMD Distance + RH Distance + MAT $Distance^2$ + MSP $Distance^2$ + CMD $Distance^2$ + RH $Distance^2$ | Seeds = MSP Distance + MAT $Distance^2$ |
| **Resistance to Herbivores** | Seeds = Leaf Herbivory + *P. spumarius* + Leaf $Herbivory^2$ + *P. spumarius*$^2$ | Seeds = *P. spumarius* + *P. spumarius*$^2$ |
| | Seeds = *S. florida* + *M. brevivitella* | Same equation |
| **Phenology** | Seeds = Flowering Date + Bolt Date + Growth Rate + Flowering $Date^2$ + Bolt $Date^2$ + Growth $Rate^2$ | Seeds = Flowering Date + Bolt Date + Growth Rate + Flowering $Date^2$ + Growth $Rate^2$ |
| **Morphology** | Seeds = SLA + Water Content + Trichome Number + $SLA^2$ + Water $Content^2$ + Trichome $Number^2$ | Seeds = SLA + Water Content + Trichome Number + Water $Content^2$ + Trichome $Number^2$ |
| **Total Phenolics** | Seeds = Leaf Total Phenolics + Leaf Total $Phenolics^2$ | Same equation |
| | Seeds = Flower Total Phenolics + Flower Total $Phenolics^2$ | Same equation |
| | Seeds = Fruit Total Phenolics + Fruit Total $Phenolics^2$ | Same equation |
| **Ellagitannins** | Seeds = Leaf Oenothein A + Leaf Oenothein $A^2$ | Same equation |
| | Seeds = Flower Oenothein A + Flower Oenothein $A^2$ | Same equation |
| | Seeds = Fruit Oenothein A | n/a |

Table S3. P-values and model coefficients for geographic, climatic, climate distance, resistance to herbivores, phenological, and morphological models predicting seed number. All variables within horizontal lines are within the same model. Bold indicates P < 0.05; underlined indicates P < 0.10.

| | Coefficient | P-value |
|---|---|---|
| **Geography** | | |
| Latitude | 13089.63 | **0.001** |
| Latitude$^2$ | -156.41 | **0.001** |
| Distance to Common Garden | -0.0003 | **0.03** |
| **Climate** | | |
| Mean Annual Temperature | 0.50 | **0.004** |
| Mean Annual Temperature$^2$ | -0.03 | **0.001** |
| Relative Humidity | 0.95 | 0.057 |
| Relative Humidity$^2$ | -0.01 | 0.055 |
| **Climate Distance** | | |
| Mean Summer Precipitation Distance | 0.002 | 0.088 |
| Mean Annual Temperature Distance$^2$ | -0.03 | **<0.001** |
| **Resistance to Herbivores** | | |
| *P. spumarius Number* | 0.29 | **<0.001** |
| *P. spumarius Number*$^2$ | -0.01 | **0.01** |
| Fruits Impacted by *S. florida* | 0.02 | **0.003** |
| *Fruits Impacted by M. brevivitella* | -0.05 | 0.058 |
| **Phenology** | | |
| Flowering Date | 0.28 | **<0.001** |

| | | |
|---|---|---|
| Flowering Date$^2$ | -0.001 | **<0.001** |
| Bolt Date | -0.04 | **<0.001** |
| Growth Rate | 65.35 | **0.049** |
| Growth Rate$^2$ | -318.70 | <u>0.071</u> |
| **Morphology** | | |
| SLA | -0.02 | 0.11 |
| Water Content | 2.38 | **0.029** |
| Water Content$^2$ | -0.02 | **0.032** |
| Trichome Number | 0.04 | **<0.001** |
| Trichome Number$^2$ | -0.0002 | **<0.001** |

Table S4. P-values and model coefficients for chemical defenses and principal components predicting seed number**.** Bold indicates P < 0.05; underlined indicates P < 0.10.

| | Coefficient | P-value |
|---|---|---|
| **Total Phenolics** | | |
| Leaf Total Phenolics | 0.02 | 0.069 |
| Flower Total Phenolics | 0.07 | **0.045** |
| Flower Total Phenolics$^2$ | -0.001 | **0.02** |
| Fruit Total Phenolics | 0.04 | **0.045** |
| Fruit Total Phenolics$^2$ | -0.0001 | 0.12 |
| **Ellagitannins** | | |
| Leaf Oenothein A | 0.02 | 0.44 |
| Leaf Oenothein A$^2$ | -0.0001 | 0.77 |
| Flower Oenothein A | 0.06 | **0.01** |
| Flower Oenothein A$^2$ | -0.002 | **0.01** |
| Fruit Oenothein A | 0.01 | **0.006** |
| **Principal Components** | | |
| PC1 | -0.09 | **0.02** |
| PC2 | 0.2 | **0.004** |

Table S5. Proportion of variance explained by all 12 principal component axes of *O. biennis* trait PCA.

| | **PC1** | **PC2** | **PC3** | **PC4** | **PC5** | **PC6** |
|---|---|---|---|---|---|---|
| Standard deviation | 2.01 | 1.29 | 1.23 | 1.05 | 0.96 | 0.86 |
| Proportion of Variance | 0.34 | 0.14 | 0.13 | 0.09 | 0.08 | 0.06 |
| Cumulative Proportion | 0.34 | 0.47 | 0.60 | 0.69 | 0.77 | 0.83 |

| | **PC7** | **PC8** | **PC9** | **PC10** | **PC11** | **PC12** |
|---|---|---|---|---|---|---|
| Standard deviation | 0.76 | 0.70 | 0.64 | 0.58 | 0.37 | 0.27 |
| Proportion of Variance | 0.05 | 0.04 | 0.03 | 0.03 | 0.01 | 0.01 |
| Cumulative Proportion | 0.88 | 0.92 | 0.95 | 0.98 | 0.99 | 1.00 |

Table S6. Loadings of first two PC axes for all traits in the *O. biennis* trait PCA.

| **Trait** | **PC1** | **PC2** |
| --- | --- | --- |
| Flowering Date | -0.36 | -0.09 |
| Water Content | 0.05 | -0.53 |
| Growth Rate | -0.05 | -0.50 |
| Bolt Date | -0.20 | 0.18 |
| SLA | -0.09 | -0.37 |
| Number of Trichomes | -0.07 | 0.14 |
| Leaf Total Phenolics | -0.23 | -0.15 |
| Flower Total Phenolics | -0.23 | 0.40 |
| Fruit Total Phenolics | -0.41 | -0.10 |
| Leaf Oenothein A | -0.38 | -0.16 |
| Flower Oenothein A | -0.42 | 0.22 |
| Fruit Oenothein A | -0.46 | -0.07 |
